# Air pollution modifies key colonization factors of the beneficial bee gut symbiont *Snodgrassella alvi* and disrupts the bumblebee (*Bombus terrestris*) gut microbiome

**DOI:** 10.1101/2023.08.04.551991

**Authors:** Hannah R. Sampson, Natalie Allcock, Eamonn B. Mallon, Julian M. Ketley, Julie A. Morrissey

## Abstract

Air pollution is the world’s largest environmental health risk. Particulate pollutants, a major component of air pollution, are detrimental to human health and a significant risk to wildlife and ecosystems globally. Black carbon, a by-product of fossil fuel and biomass burning, is a key constituent of air pollution with levels continuing to increase worldwide. Here we describe the effects of black carbon on the beneficial gut microbiome of an important global insect pollinator, the buff-tailed bumblebee (*Bombus terrestris*). Our data shows that exposure to black carbon particulates alters the biofilm structure, gene expression and initial adhesion of beneficial bee gut coloniser, *Snodgrassella alvi in vitro.* Additionally, our results show that black carbon disrupts adult *Bombus terrestris* gut microbiome composition, a vital component to bee health. Exposure to black carbon increased bees’ viable gut bacteria and significantly altered the abundance of beneficial core bacteria *Gilliamella* and *Bombilactobacillus* in the microbiome. These findings demonstrate that exposure to black carbon air pollution has direct, measurable effects on bees’ beneficial gut commensal bacteria and microbiome. Together these data highlight that particulate pollutants are an underexplored risk for the health of insect pollinators.

## Introduction

Air pollution is the world’s largest environmental health risk, with around 90% of the global population residing in areas that exceed the World Health Organisation’s air quality guidelines (WHO, 2018). Air pollutants originating from both natural and anthropogenic sources contaminate our atmosphere and can be transported large distances, depositing on plants, soil and water sources. Air pollution is comprised of gaseous components and solid particulate matter (USEPA, 2016b). Black carbon is a major component of particulate matter and is produced from the incomplete combustion of fossil fuels and biomass, with levels continuing to rise due to increasing urbanisation, industry and wildfires (Ditas *et al*., 2018). Particulate matter exposure is associated with increased human respiratory and cardiovascular disease severity (Cohen *et al*., 2017), and in the environment exposure alters ecosystem diversity, nutrient availability and is detrimental to wildlife health (USEPA, 2016a; Li *et al*., 2021).

Both honeybees and bumblebees are vital to the maintenance of natural ecosystems and are essential pollinators in agriculture (Klein *et al*., 2007; Potts *et al*., 2016; Cameron and Sadd, 2020) making the study and maintenance of their health a crucial area of research. Bees house a distinct, specialised community of core gut bacterial symbionts (the bee gut microbiome) that are unique to the bee gut and colony environment (Koch and Schmid-Hempel, 2011b, 2011a; Martinson *et al*., 2011; Engel *et al*., 2012; Moran *et al*., 2012; Anderson *et al*., 2013; Kwong *et al*., 2017). The maintenance and balance of this core gut microbial community is a key element to bee health assisting in the digestion of essential nutrients, promoting host weight gain and providing protection from pathogens and environmental stressors (Koch and Schmid-Hempel, 2011b; Kesnerova *et al*., 2017; Zheng *et al*., 2017; Motta *et al*., 2018).

Core bacterial phylotypes (*Snodgrassella, Gilliamella, Bifidobacteriaceae, Lactobacillus* Firm-4 and *Lactobacillus* Firm-5) have been identified in almost every corbiculate eusocial bee (e.g. honeybees and bumblebees) and usually make up the majority of their gut microbiome (Koch and Schmid-Hempel, 2011a; Martinson *et al*., 2011, 2012; Moran *et al*., 2012; Cariveau *et al*., 2014; Meeus *et al*., 2015; Kwong *et al*., 2017). *Lactobacillus* Firm-4 (*Bombilactobacillus bombi*) and *Lactobacillus* Firm-5 reside in the hindgut (ileum and rectum) and play a key role metabolising pollen, providing nutrients for the host (Kesnerova *et al*., 2017; Zheng *et al*., 2017; Zijing Zhang *et al*., 2022). *Snodgrassella* and *Gilliamella* predominantly reside on the ileum epithelium in a biofilm structure and engage in syntrophic cross-feeding of nutrients (Martinson *et al*., 2012; Kwong *et al*., 2014; Kesnerova *et al*., 2017; Hammer *et al*., 2021). *Snodgrassella* is a key gut coloniser, able to persist in an otherwise sterile gut without other members of the gut microbiome (Kwong *et al*., 2014). Several genes are essential to this colonisation process including adhesins and type IV pilus genes (Kwong *et al*., 2014; Powell *et al*., 2016; Leonard *et al*., 2018). Core bacteria interact with the bee host influencing their immune response and hormone signalling, demonstrating that core symbionts not only breakdown food but also shape the internal physiology of the bee (Kesnerova *et al*., 2017; Zheng *et al*., 2017; Horak *et al*., 2020).

Bumblebees hatch with few to no bacteria in their intestine and obtain their characteristic mature adult gut microbiome over four to six days through interactions with nestmates and hive material in their colony (Koch and Schmid-Hempel, 2011b; Meeus *et al*., 2013; Billiet *et al*., 2017). Successful establishment of the microbiota plays a major role in protecting bees against pathogens (Koch and Schmid-Hempel, 2011b). Bees taken away from the colony early, before their mature gut microbiome has been established, do not develop a normal gut microbiome composition and have increased infection levels (Koch and Schmid-Hempel, 2011b; Billiet *et al*., 2017).

Once mature, the indoor-reared bumblebee microbiome composition remains stable as they age (Parmentier *et al*., 2016), but this mature microbiome can be perturbed by environmental factors, for example exposure to environmental metal pollutants (Rothman *et al*., 2019). There is a wealth of literature concerning pollinator health and pesticides, but substantially less research on air pollution and insects (Feldhaar and Otti, 2020), highlighting the need for a better understanding of how pollutants affect insect microbiomes and health.

Bees have long been used as indicator organisms to assess the impact and prevalence of atmospheric pollutant contamination (Bromenshenk *et al*., 1985). Bees accumulate atmospheric particulates on their head, antennae, legs and wings (Negri *et al*., 2015; Thimmegowda *et al*., 2020) and particulates also contaminate collected pollen (Papa *et al*., 2021). This means that pollutants are not only in contact with bees during foraging but are present in their food stores. *Apis dorsata* (Giant Honeybee) collected from highly polluted sites were associated with significant changes to flower visitation, heart rate and survival (Thimmegowda *et al*., 2020), indicating the potential systemic impact pollution has on bees, their behaviour and longevity. Understanding more about how bees are affected by particulate pollutants is important to determine the full impact on bee health.

Previous studies have shown that exposure to the major particulate air pollutant black carbon changes the behaviour of human pathogens *Staphylococcus aureus* and *Streptococcus pneumoniae* altering their biofilm formation *in vitro* and in murine respiratory tract colonisation (Hussey *et al*., 2017; Purves *et al*., 2022). Pre-exposure to black carbon prior to inoculation altered virulence gene expression and colonisation, demonstrating that *S. aureus* genetically adapts to black carbon exposure (Purves *et al*., 2022). The discovery that black carbon can alter bacterial biofilm structure and bacterial colonisation *in vivo* showed that particulate pollutants, such as black carbon, could have more, currently unknown, influences on microbial life.

Here we report that not only does air pollution affect the behaviour of bee gut commensals, but also perturbs the core bee gut microbiome. We investigated the impact of black carbon on model bee gut bacteria *Snodgrassella alvi*, a prominent member of the core bee gut microbiome, finding *S. alvi* adhesion and *in vitro* biofilm formation was altered by black carbon. Importantly, exposure of black carbon to the native UK pollinator the buff-tailed bumblebee (*Bombus terrestris*), reared in controlled laboratory conditions, caused direct, measurable effects on the mature bee gut microbiome composition. These findings have noteworthy implications for the impact of air pollution on bee health.

## Experimental procedures

### Bacterial growth conditions

The bee gut commensal type strain *Snodgrassella alvi* wkB2 (Kwong and Moran, 2013) (DSMZ, DSM 104735) was used in this study. *S. alvi* was stored at −80°C and cultivated from frozen glycerol stocks on blood agar. Unless otherwise stated, bacteria were grown using blood agar supplemented with 5% horse blood (Oxoid), Brain Heart Infusion broth (BHI, Oxoid) and incubated statically in 5% CO_2_ and 37°C conditions.

### Black carbon and quartz particles

Black carbon (BC) (Sigma-Aldrich 699632) powder was suspended in sterile ddH_2_O (Hussey *et al*., 2017; Purves *et al*., 2022) to make a stock solution at 2-10 mg/mL. Quartz particles (Distrilab BCR66) were also suspended in sterile ddH_2_O for use in this work. Black carbon concentrations used in this study were based on previous research into the effect of black carbon on the host and host-associated bacteria, using high concentration suspensions to model long-term environmental exposure (Hussey *et al*., 2017; Purves *et al*., 2022). Particle sizes used in this study (BC <0.5 µm and quartz 3.5 – 0.35 µm) are within the range of environmental particulate pollutant sizes found to accumulate on wild bees (10 – 0.02 µm) (Thimmegowda *et al*., 2020; Capitani *et al*., 2021).

### Pre-treated media

Sterile BHI media supplemented with 100 µg/mL black carbon was vortexed for 3 seconds and incubated at 37°C overnight. Media was centrifuged for 10 minutes at 3220 x g 22°C and supernatant was filtered through a 0.2 µm filter to remove any remaining particulates.

### Snodgrassella alvi biofilm assay

Overnight cultures of *S. alvi* wkB2 were set to an optical density of 0.02 at 600 nm in BHI, aliquoted into a 12 well plate with or without 100 µg/mL of black carbon and incubated in 5% CO_2_ at 37°C for 2 hours, 8 hours, 24 hours or 48 hours. The assay protocol from Hussey *et al*. (2017) was followed except for the use of BHI in place of PBS for washes and dilutions. Fractions were vortexed for 10 seconds, samples were serially diluted in BHI and plated onto blood agar plates to count colony forming units (CFU). To image biofilms, samples were set up and incubated as above, supernatant was removed, and the biofilm stained with crystal violet for 5 minutes. The crystal violet stain was removed, biofilms were washed with ddH_2_O and imaged using Immunospot® (Cellular Technology Limited).

### Scanning electron microscopy

Bacterial cultures were inoculated onto sterile plastic coverslips (Agar Scientific AGL4193) within 12 well plates and incubated at 5% CO_2_ and 37°C for 24 hours. After incubation, coverslips were fixed with 2.5% glutaraldehyde in 0.1 M Sörensens buffer for 2 hours at room temperature (Dykstra and Reuss, 2003). After fixing, biofilms were washed in PBS, ddH_2_O and dehydrated in an ethanol series, then a graded series of ethanol:hexamethyldisilazane mixtures. Samples were air dried overnight, mounted onto pin stubs, gold coated and viewed with a Hitachi S-3000H. To image biofilm structures, stubs were tilted 90° and biofilm protrusions were imaged in cross sections by random selection, framing and focussing then protrusion areas were measured from the surface of the coverslip and analysed using the image threshold function in ImageJ (Rasband, 2018).

### RNA extraction

To prepare *S. alvi* planktonic growth samples for RNA extraction *S. alvi* was grown to exponential phase (OD_600 nm_ = 0.08) in BHI with and without 100 µg/mL black carbon. RNA was stabilised with the addition of one fifth the culture volume of 1:19 Phenol:Ethanol and incubated on ice for ∼25 minutes. The culture was centrifuged at 4,000 rpm (3220 x g), 4°C for 5 minutes to pellet the cells. The supernatant was discarded, and the pelleted cells were stored at −80°C.

To prepare *S. alvi* biofilm samples for RNA extraction *S. alvi* biofilms were grown in BHI with and without 100 µg/mL black carbon and incubated for 24 hours. Supernatant and wash fractions were removed and discarded, RNA was stabilised with the addition 1 mL RNAlater (ThermoFisher) and incubated for 5 minutes at room temperature. The biofilm fraction was transferred to a microcentrifuge tube, centrifuged (11400 x g) for 5 minutes. The supernatant was discarded, and the pelleted cells were stored at −80°C.

RNA was extracted following a similar protocol as previous work (Purves *et al*., 2022), except frozen pellets were thawed and lysed in 200 µL of Tris-EDTA buffer containing 20 µL of lysosyme (1 mg/mL) and incubated at 37°C for 15 minutes prior to being mechanically disrupted using Lysing Matrix B tubes and MP Biomedicals FastPrep-24 Classic instrument (MP Biosystems). RNA was extracted from the supernatant using a Direct-zol RNA Miniprep kit (Zymogen) following the manufacturer’s instructions and samples were further treated as described in Purves *et al.,* (2022).

### Quantitative reverse transcriptase PCR (qRT-PCR)

Total RNA was converted to cDNA using Superscript IV VILO Master Mix reverse transcriptase (Invitrogen) and 1 ng of cDNA was used for each qPCR reaction. Reactions (10 µL) were carried out in triplicate using SYBR Green Master mix in a 7300 Fast System (Applied Biosystems) following manufactures instructions. Relative gene expression for each sample (primer details Table S1) was normalised to the expression of endogenous control 16S rRNA gene and analysed relative to *S. alvi* cultured without black carbon. Relative quantification was calculated using ΔΔCt analysis (Livak and Schmittgen, 2001).

### Bumblebee rearing and husbandry

Four *Bombus terrestris audax* colonies, were ordered from Biobest (Westerlo, Belgium) in January 2019. Colonies were reared in constant red-light conditions at 28°C and 60% humidity. They were fed 60% v/v apiary solution (Meliose-Roquette, France) and pollen (Percie du sert, France) *ad libitum*. Callow workers, less than 24 hours old, were caught, labelled on the thorax with a marker, their number recorded and returned to the colony. Once they had obtained their mature, adult gut microbiome (7± 1 day old (Meeus *et al*., 2013)), marked bees were placed into rearing boxes of four to six individuals and designated either treatment or control group. Experimental groups were reared in constant red-light conditions at 28°C and 60% humidity and fed pollen and apiary solution (70% v/v) *ad libitum*.

### Experimental setup, black carbon exposure and sampling

Both experimental groups were fed apiary solution (70% v/v) for two days. After faecal sample collection on day two the control group was provided fresh apiary solution and the treatment group was given apiary solution containing black carbon (0.495 mg/g black carbon as calculated in Figure S1) for the remaining four days of the experiment. Black carbon concentration was calculated based on concentrations used in previous animal work (Hussey *et al*., 2017) and accounted for average daily consumption of apiary solution (Tyler *et al*., 2006) and weight (Bumblebee.org, 2017) (Figure S1).

### Bee survival and activity monitoring

Activity was recorded from one bee, selected at random (using a random number generator) per experimental box. Twice each day bees were randomly selected per experimental box and their active time was recorded for 150 seconds. Survival was measured periodically throughout the experiment to monitor host mortality.

### Faecal sampling, DNA extraction and total 16S rRNA copy number

Bees were individually caught, placed in ventilated universal tubes and anaesthetised on ice for ∼20 minutes prior to sample collection. On experimental days two and six faecal samples were taken from anaesthetised bees by gently squeezing the abdomen (Bailey *et al*., 1983). Fresh faecal samples were diluted in 100 µL of sodium phosphate buffer from the FastDNA SPIN kit for soil (Qbiogene, Carlsbad, 2019) and briefly vortexed. From each sample, 20 µL was taken for bacterial culture and the remaining sample was frozen at −20°C. For bacterial culture, each sample was serially diluted, plated onto De Man, Rogosa and Sharpe (MRS) agar and blood agar and incubated in microaerobic (34°C, 10% CO_2_, 85% N_2_, 5% O_2_) conditions for 72 hours. After incubation, viable counts were recorded to calculate CFU.

Faecal samples were thawed, pooled according to experimental box and processed with the FastDNA SPIN kit for soil (Qbiogene, Carlsbad, 2019). Samples were paired-end sequenced on an Illumina 1.9 platform. The V3-V4 region of the 16S rRNA gene was amplified with primers 341F (5-CCTAYGGGRBGCASCAG-3) and 806R (5-GGACTACNNGGGTATCTAAT-3). Negative controls for the DNA extraction kit were amplified for the V4 region with primers 515F (5-GTGCCAGCMGCCGCGGTAA-3) and 806R (5-GGACTACHVGGGTWTCTAAT-3).

Total 16S rRNA copy number was amplified from all samples using universal bacterial primers (Table S1) as in previous studies (Raymann *et al*., 2017; Motta and Moran, 2020). Reactions (10 µL) were carried out in triplicate using SYBR Green Master mix in a 7300 Fast System (Applied Biosystems), using PCR cycles as described in Raymann *et al.,* (2017). Standard curves from the amplification of the cloned target sequence in a pGEM-T vector (Promega) were used to quantify 16S total copy number per sample.

### Bioinformatics pipeline

Individual sample quality was determined with FastQC (Andrews, 2010) and summarised with MultiQC v1.9 (Ewels *et al*., 2016). Illumina paired-end reads were processed in Qiime2 v2020.2 (Bolyen *et al*., 2019). Barcodes and primers were trimmed with DADA2 (Callahan *et al*., 2016), sequences were denoised, filtered for quality, merged, chimeric sequences were removed and features identified. The bee-associated microbial community database BEExact (Daisley and Reid, 2021), was trained for the appropriate 16S region and used to taxonomically assign samples’ amplicon sequence variants (Bokulich *et al*., 2018). Differential abundance analysis was carried out in R using ANCOM-BC (Lin and Peddada, 2020). Rarefaction, filtering and alpha and beta diversity analysis (Faith, 1992; Lozupone *et al*., 2007) was carried out in Qiime2 and R (R Core Team, 2013) utilising packages phyloseq (McMurdie and Holmes, 2013), ggplot2 (Wickham, 2009) and the tidyverse (Wickham *et al*., 2019).

### Graphical output, data and image analysis

Graphs were produced and data analysed using R statistical software (R Core Team, 2013) or GraphPad Prism (version 7.0.4 for Windows, GraphPad Software, San Diego, California USA). Scanning electron microscopy images were processed using ImageJ (Rasband, 2018).

## Results

### Black carbon increases initial adhesion of the beneficial bee gut symbiont Snodgrassella alvi, changing biofilm formation

Previous studies have shown that exposure to black carbon changes the host colonisation of human bacterial pathogens *S. aureus* and *S. pneumoniae* (Hussey *et al*., 2017; Purves *et al*., 2022). However, the direct effect of black carbon on beneficial bacterial commensals, such as bee gut commensals, has not been established. *S. alvi* is a prominent, beneficial bee gut commensal (Moran *et al*., 2012; Kwong *et al*., 2014; Powell *et al*., 2016; Horak *et al*., 2020) and is used in this study as a model bee commensal to establish whether black carbon affects core bee gut bacteria.

Black carbon concentrations used in this study were based on previous published work studying the effects of black carbon on the host and host-associated bacteria (Hussey *et al*., 2017; Purves *et al*., 2022). Dose response analysis showed no significant effect (*F*_(3, 8)_ = 1.58, *p* > 0.05) on *S. alvi* growth with higher concentrations of black carbon (Figure S2). For *in vitro* experiments, 100 µg/mL of black carbon was used because it had minimal effect on *S. alvi* growth (Figure S2, S3) and is consistent with previous studies.

*In vivo, S. alvi* predominantly colonises the ileum, binding to ileal epithelial cells in a biofilm structure (Martinson *et al*., 2012; Hammer *et al*., 2021). The effect of black carbon on *S. alvi* wkB2 biofilm formation was measured *in vitro* by quantifying the viability of different fractional components of the biofilm. *S. alvi* viability was measured in three biofilm fractions; supernatant (planktonic), wash (loosely adherent) and adherent biofilm (Hussey *et al*., 2017). *S. alvi* biofilms exposed to black carbon had significantly higher colony forming units (CFU) at 2 hours (initial adhesion) (*F*_(2, 12)_ = 20.98, *p* ˂ 0.01), 8 hours (early biofilm) (*F*_(2, 12)_ = 33.04, *p* ˂ 0.01) and 24 hours (early biofilm) (*F*_(2, 12)_ = 17.01, *p* < 0.05) compared to the control (Figure 1A, 1B, 1C). Conversely, there were significant decreases in the CFU of black carbon treated supernatant fractions at 2 hours (*p* ˂ 0.001), 8 hours (*p* ˂ 0.001) and 24 hours (*p* ˂ 0.001). At the 48 hours (established biofilm) timepoint, no black carbon treated fractions were significantly different to any control fractions (Figure 1D) (*F*_(2, 12)_ = 3.484, *p* > 0.05). When the CFU of each fraction was combined, there was no significant difference between control and black carbon treated *S. alvi* total CFU (Figure S4) (*F*_(7, 39)_ = 33.97, 2 hours *p* > 0.05, 8 hours *p* > 0.05, 24 hours *p* > 0.05, 48 hours *p* > 0.05), therefore black carbon changed *S. alvi* biofilm behaviour but not overall growth in these conditions.

**Figure 1.**
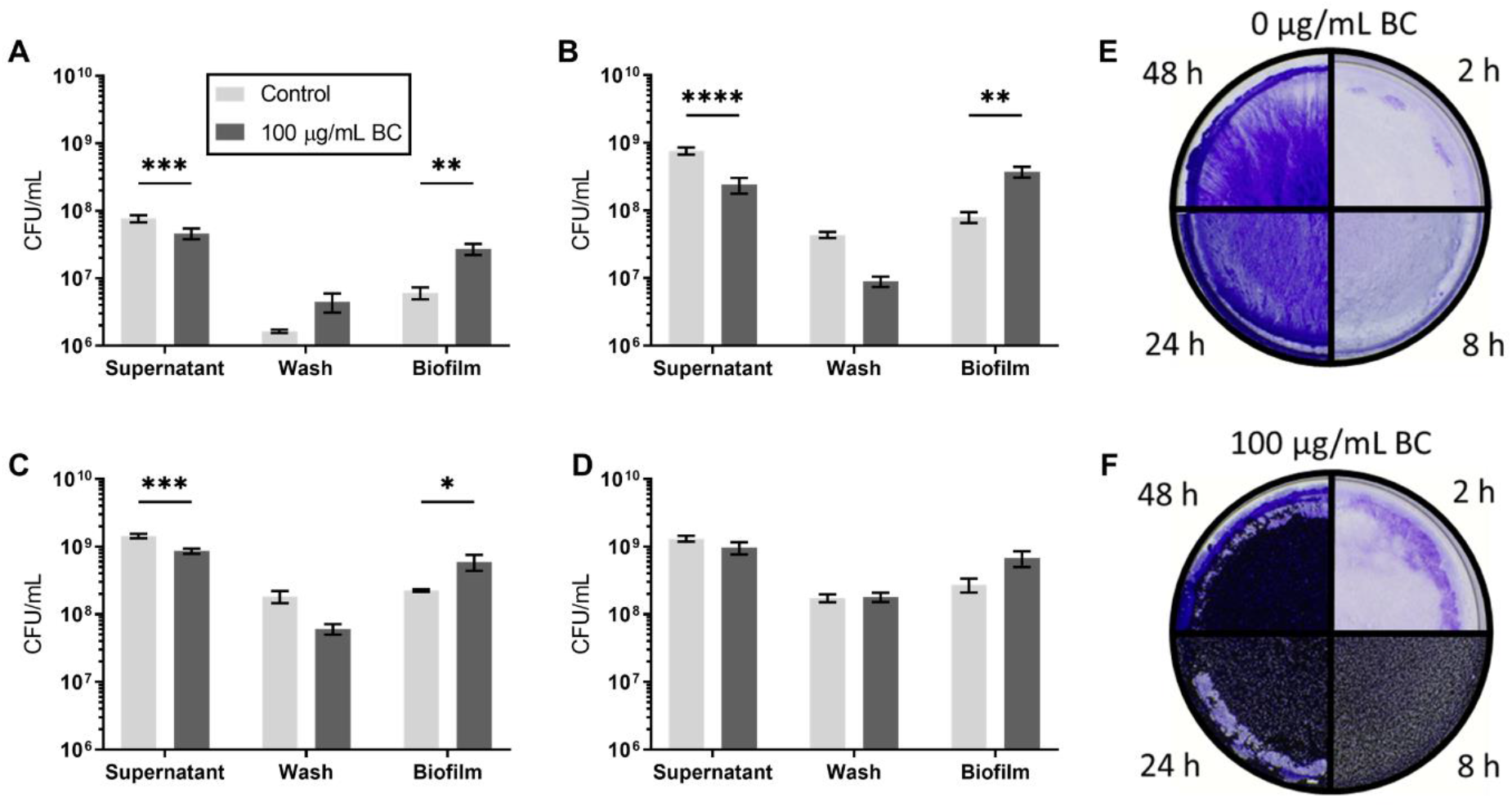
Black carbon increases *Snodgrassella alvi* early biofilm formation. *S. alvi* wkB2 was grown in BHI with 0 or 100 μg/mL black carbon (BC), aliquoted into 12 well biofilm plates and incubated at 37° C in 5% CO2 conditions for 2 hours (A), 8 hours (B), 24 hours (C) or 48 hours (D). Biofilms were separated into fractions (supernatant (planktonic growth), wash (loosely adherent) and biofilm), serial diluted and plated to determine bacterial growth. Data are presented as CFU/mL and error bars represent standard error of the mean of n=5 biological repeats. Two-way ANOVAs were performed followed by Šidák correction, significant differences (* p < 0.05, ** p < 0.01, *** p < 0.001 and **** p < 0.0001) are indicated. Images of crystal violet stained biofilms at different timepoints treated with 0 (E) or 100 μg/mL black carbon (F).

Biofilms treated with 0 or 100 µg/mL black carbon were visualised with crystal violet staining (Figure 1E and 1F). Biofilm quantification with crystal violet was impeded by black carbon but demonstrates the clear incorporation of black carbon into the biofilm from 8 hours. No black carbon was observed in the biofilm at 2 hours even though there is a significant increase in viable cells in the biofilm fraction (Figure 1A) suggesting that early adhesion is dependent on black carbon altering bacterial behaviour. Together these data show that black carbon significantly increases *S. alvi* wkB2 adherence in the early phases of biofilm formation.

### The effect of black carbon particles on Snodgrassella alvi biofilm structure

To determine if the physical nature of the black carbon signal results in a biofilm change, *S. alvi* biofilms were grown for 24 hours in BHI media that was pre-treated by adding black carbon to the media, then removing the particles by filtration prior to *S. alvi* inoculation. The effect of pre-treated media on *S. alvi* 24-hour biofilm formation was compared to black carbon treated and control biofilms. *S. alvi* biofilm formation in black carbon pre-treated media showed a similar response to the control at 24 hours, with a significantly higher supernatant CFU (*F*_(4, 36)_ = 8.28, *p* ˂ 0.001) in both control and pre-treated media conditions compared to the black carbon-treated condition, but a non-significant change in biofilm CFU (*p* > 0.05) (Figure 2A). Only *S. alvi* biofilms grown in the presence of black carbon particles had a significantly altered biofilm structure (*p* < 0.05) therefore, the physical presence of black carbon particles is required for these *S. alvi* biofilm alterations.

**Figure 2.**
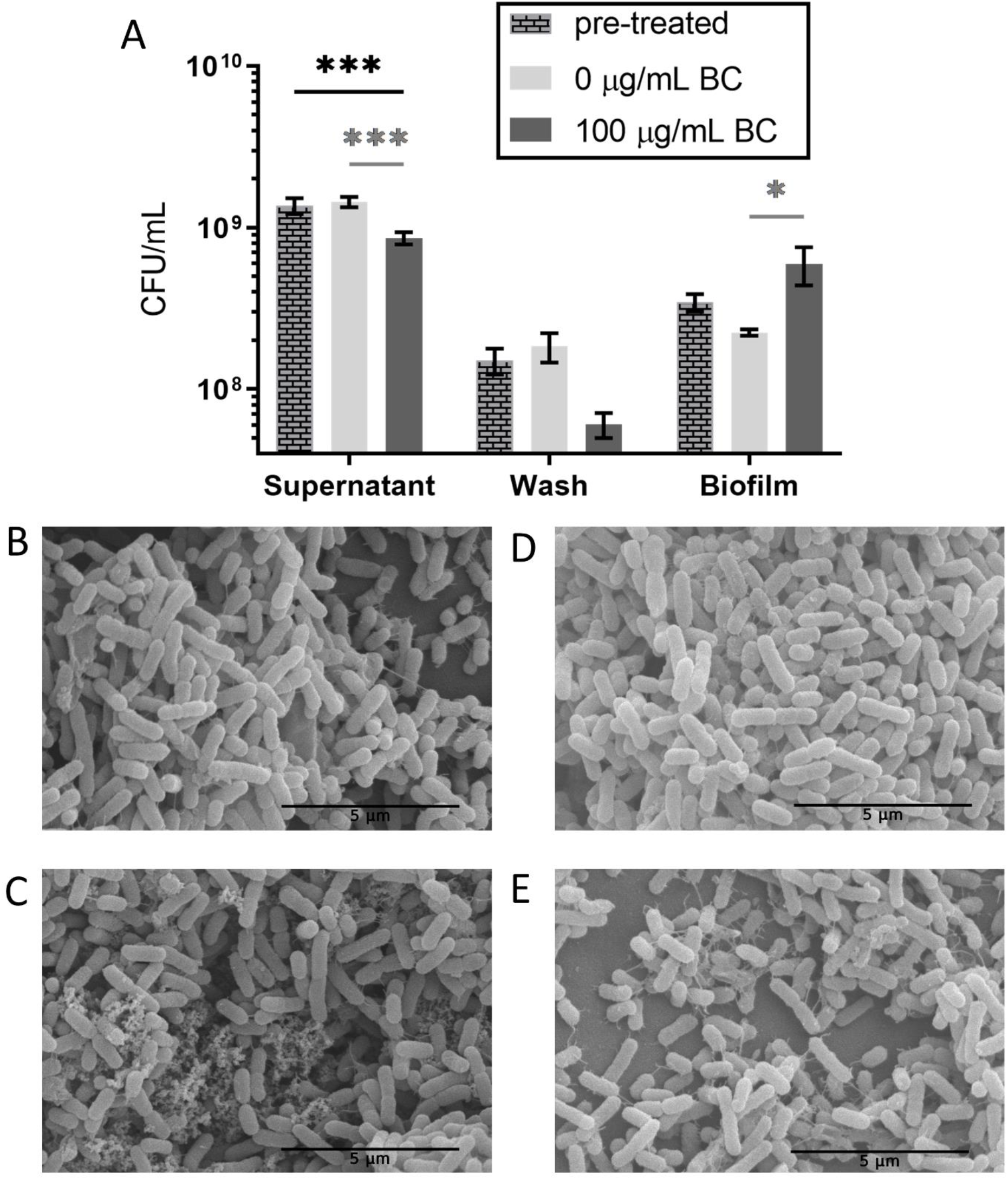
Black carbon particles change *Snodgrassella alvi* biofilm structure. A) Pre-treated CFUs are presented alongside 0 or 100 μg/mL black carbon (BC) 24 hour biofilms, biological repeats n=5. Error bars represent standard error of the mean. Two-way ANOVA with Šidák correction was performed and significant differences (* p < 0.05 and *** p < 0.001) are indicated in black between the pre-treated condition and 0 or 100 μg/mL black carbon, and in grey between 0 or 100 μg/mL black carbon conditions. Scanning electron microscopy images of *S. alvi* wkB2 biofilms grown in BHI with 0 μg/mL black carbon (B), 100 μg/mL black carbon (C), pre-treated media (D) or 100 μg/mL quartz (E).

To establish what architectural and structural biofilm changes occur in the presence of black carbon, scanning electron microscopy was conducted on *S. alvi* biofilms. Biofilms were grown for 24 hours, fixed, dehydrated through ethanol and hexamethyldisilazane and imaged. *S. alvi* biofilms were grown in 0 μg/mL black carbon, 100 μg/mL black carbon or pre-treated media. Another condition, 100 μg/mL quartz, was used as a control to determine if particles of a similar size to black carbon, but chemically inert, effected *S. alvi* biofilm structure.

In all conditions, higher magnification images show hair-like projections (Figure 2B, 2C, 2D, 2E) and black carbon can be seen incorporated into the biofilm structure in close proximity to *S. alvi* cells (Figure 2C). Pre-treated and quartz-treated biofilms (Figure 2D, 2E) appear thinner, with more gaps, similar to the 0 μg/mL black carbon treated samples (Figure 2B). Only black carbon treated biofilms (Figure 2C) showed an altered structure, which agrees with CFU biofilm measurements.

### Black carbon changes Snodgrassella alvi gene expression

Previous work showed that exposure of *S. aureus* to black carbon particles changed *S. aureus* global gene expression (Purves *et al*., 2022). To determine whether black carbon changes *S. alvi* behaviour through altered gene expression, RNA was extracted from *S. alvi* exponential planktonic growth and *S. alvi* 24-hour biofilms. The expression of genes potentially involved in biofilm structure (*pilD*) and regulation (*lysR*) was assessed with qPCR. The *pilD* gene encodes a prepilin peptidase; an essential gene for *S. alvi* adherence, colonisation and biofilm formation *in vivo* and *in vitro* (Powell *et al*., 2016). In other bacterial species the transcriptional activator *lysR* is important for regulating biofilm formation and colonisation (Seib *et al*., 2007; Maddocks and Oyston, 2008). *S. alvi* possesses a homologue of *lysR* (Rothman *et al*., 2020) but its function is unconfirmed.

*S. alvi* biofilms formed in the presence of black carbon had no significant change in *pilD* expression (*F*_(1, 8)_ = 7.09, *p* > 0.05) but showed a significant decrease in *lysR* expression (*p* < 0.05) (Figure 4). These changes were not observed in exponential planktonically grown cells (Figure 4) (*F*_(1, 8)_ = 0.11, *p* > 0.05).

**Figure 3.**
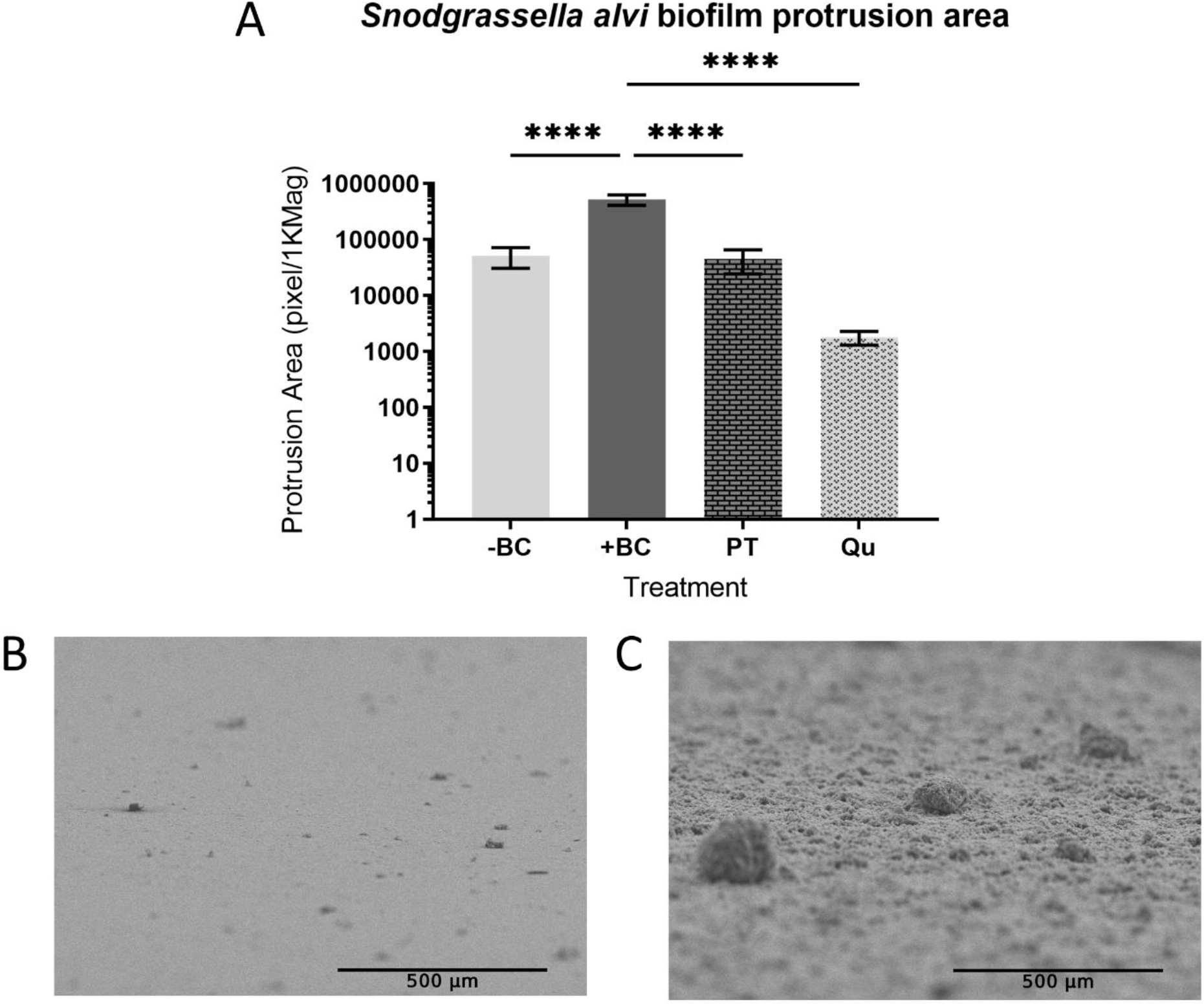
Black carbon particles change *Snodgrassella alvi* biofilm architecture. *S. alvi* wkB2 biofilms were grown for 24 hours in BHI with 0 μg/mL black carbon (-BC), 100 μg/mL black carbon (+BC), 100 μg/mL quartz (Qu) or in black carbon (100 μg/mL) pre-treated BHI (PT). Biofilms were processed and protrusions imaged with scanning electron microscopy. A) *S. alvi* biofilm protrusion areas were normalised to 1,000x magnification and analysed with a one-way ANOVA and Tukey’s multiple comparisons test, significant differences (**** p < 0.0001) are shown, n=6 images for each condition. Representative scanning electron microscopy images of *S. alvi* biofilm protrusions treated with 0 μg/mL black carbon (B) or 100 μg/mL black carbon (C).

**Figure 4.**
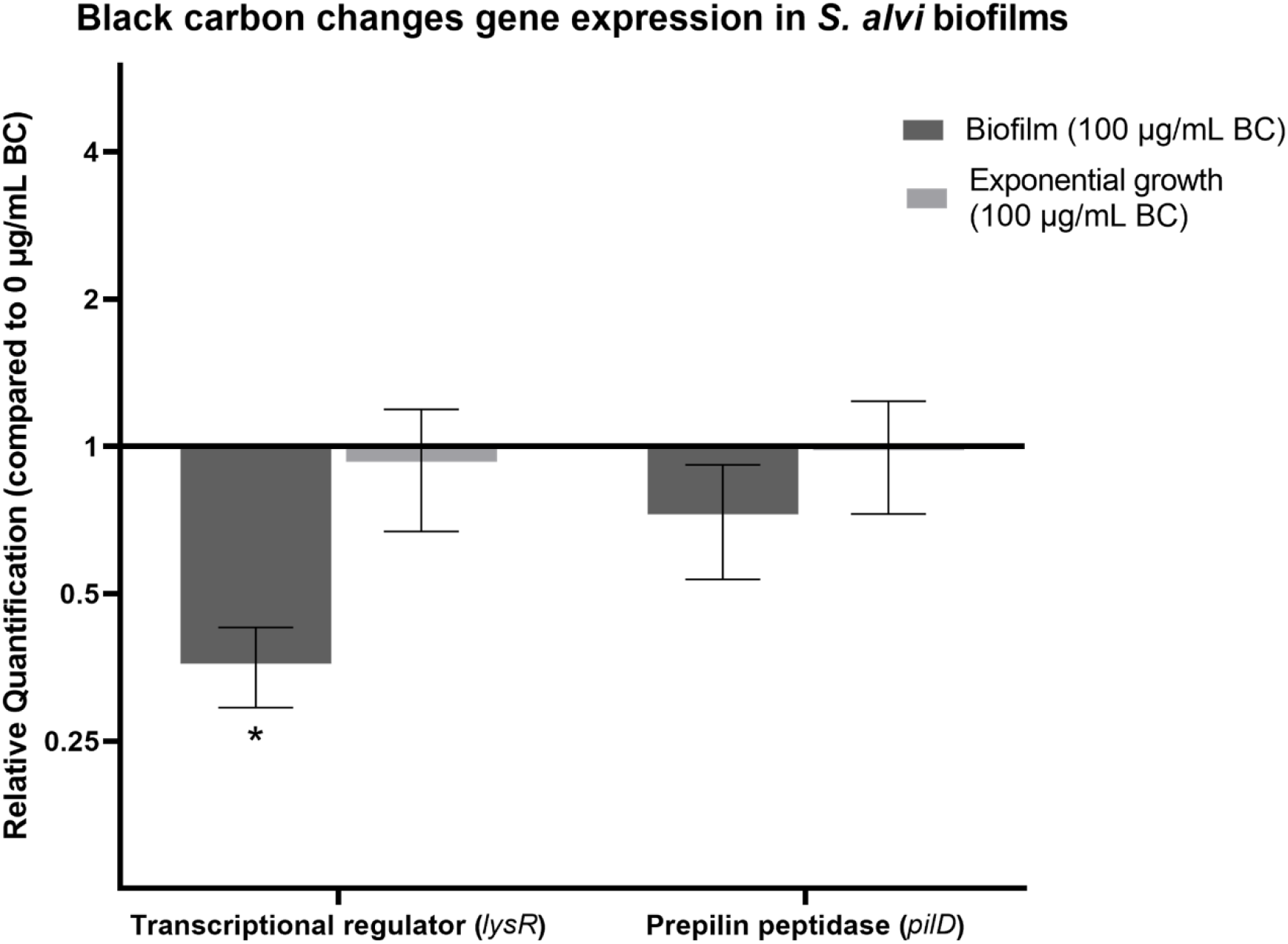
The effect of black carbon on *Snodgrassella alvi* gene expression. Relative fold change of *S. alvi* wkB2 *lysR* and *pilD* gene expression when grown exponentially or in a biofilm with 100 μg/mL black carbon (BC). Relative quantification is the fold change in expression relative to 0 μg/mL black carbon. The endogenous control, *16S* rRNA, was used as its expression level did not change between control and black carbon treatment. Biological replicates n=3, error bars represent standard error of the mean. A two-way ANOVA was performed on each sample type (exponential planktonic growth and biofilm) with Fisher’s LSD correction, significant differences (* p < 0.05) are indicated.

### Black carbon perturbs the bumblebee gut microbiome at a concentration non-toxic to the host

Given that black carbon changes the behaviour of bee gut commensal *S. alvi in vitro,* there is potential for black carbon to impact the bee gut microbiome. The impact of black carbon exposure on the bumblebee gut microbiome and subsequent changes to bee activity and survival were investigated in lab-reared adult *B. terrestris*.

Age controlled adult worker bees with a fully formed, mature gut microbiome (7±1 day old) were moved into experimental boxes and designated either treatment or control group. Both experimental groups were fed apiary solution for two days (Pre), after which fresh apiary solution was provided alone (control) or with 0.495 mg/g black carbon (treatment) (Post) (Figure 5A).

**Figure 5.**
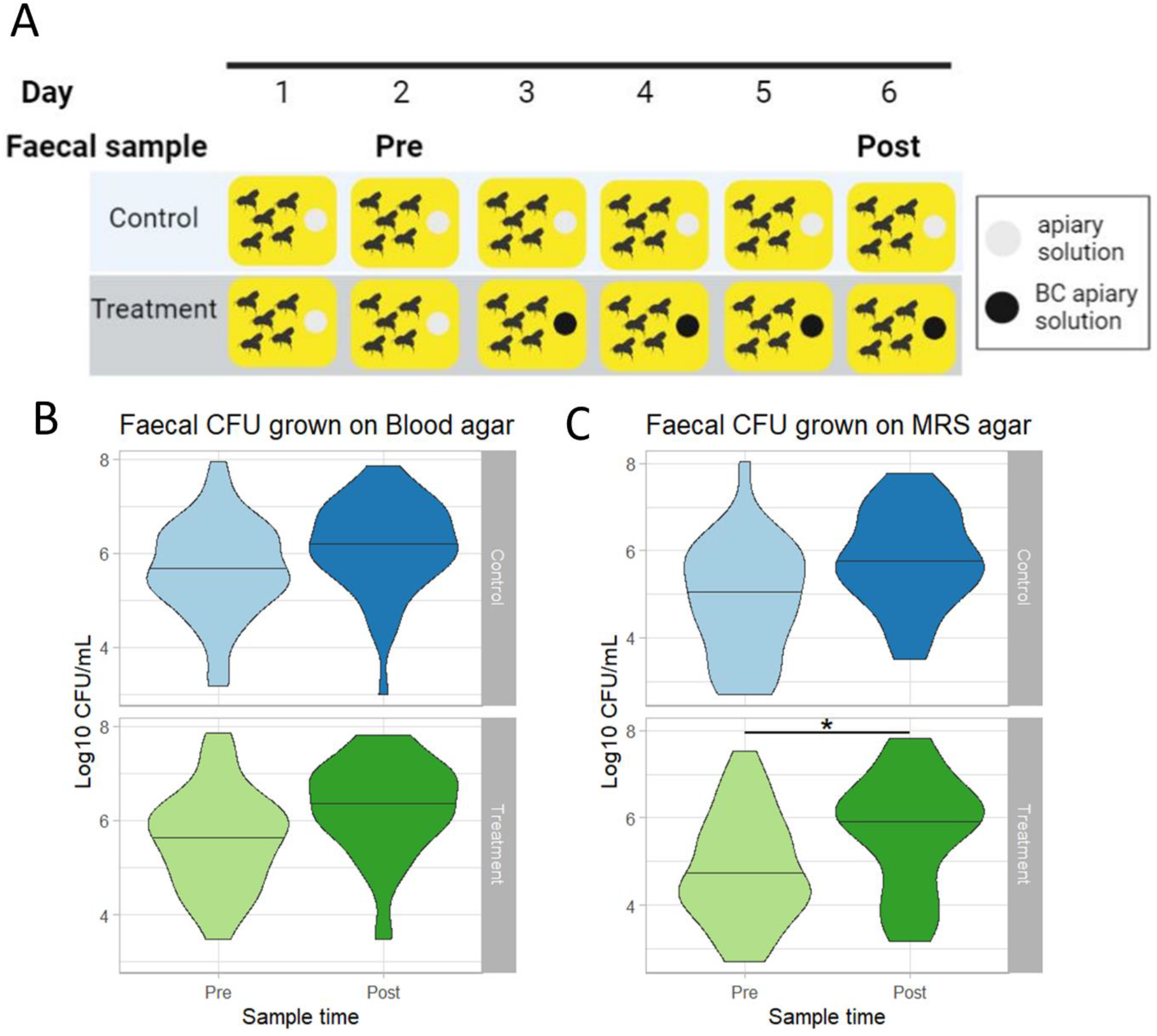
The effect of black carbon on adult *Bombus terrestris* viable gut bacteria. A) Experimental setup showing feeding patterns with apiary solution or black carbon (BC) laced apiary solution (0.495 mg/g) and sample collection across a daily period (created with Biorender.com). B-C) *B. terrestris* faecal CFU per experimental group (control n=42; black carbon treatment n=52), lines represent median. Faecal samples were grown on blood agar (B) or MRS agar (C) and incubated for 72 hours in microaerobic conditions (34°C, CO2: 10%, O2:5%, N2:85%). CFU results from experimental groups were analysed with a general linear model and a two-way ANOVA with multivariate p value adjustment, significant differences (* p < 0.05) are indicated.

To determine if black carbon influenced bee behaviour, bee active time was recorded for control and treatment experimental groups Pre and Post treatment. Bees were randomly selected per experimental box and their activity was recorded for 150 seconds. There was no significant difference in active time Pre and Post for either experimental group (Figure S5) (control W = 100.5, *p* > 0.05; treatment W = 86.5, *p* > 0.05). The effect of black carbon treatment on bee mortality rates was assessed by monitoring bee survival periodically throughout the experiment. Control and treatment groups had no significant difference (W=14.5, *p* > 0.05) in mortality with ∼80% of bees from each experimental group surviving until the end of the experiment (Figure S6). As bees treated with black carbon did not have any significant differences in survival or activity levels compared to controls the dose of black carbon used does not appear to be overtly toxic to the host.

To determine the effects of black carbon on gut microbiome composition, faecal samples were taken (Bailey *et al*., 1983) at two timepoints Pre (day two) and Post black carbon exposure (day six). Faecal samples were used to monitor the gut microbiome composition over time. Part of each sample was cultured to quantify viable bacteria; the remaining samples were pooled by experimental box and DNA extracted for 16S rRNA amplicon sequencing.

For quantification of viable bacteria, a complex non-selective medium (blood agar) and a lactobacilli-selective culture medium (MRS agar) were used to culture bee faecal samples taken at Pre and Post timepoints from control bees and those treated with black carbon. Bacterial viability of samples grown on blood agar was not significantly different between experimental groups or sample time (χ^2^(1) = 352.79, control *p* > 0.05; treatment *p* > 0.05) (Figure 5B). Lactobacilli, a subset of the bee gut microbiome selectively cultured on MRS, showed no significant difference (*p* > 0.05) in the control group bacterial viability. However, there was a significant increase (*p* < 0.05) in CFU in the treatment group Post black carbon treatment compared to Pre-treatment (Figure 5C). Black carbon exposure changes the adult bee gut microbiome as MRS-culturable bacteria were significantly increased after black carbon treatment.

To determine which members of the gut microbiome are altered with black carbon treatment, the remaining faecal samples were pooled by experiment box and DNA extracted. Total 16S rRNA gene copy number was quantified by qPCR, finding no significant difference (*F*_(3, 18)_ = 0.86, *p* > 0.05) in absolute 16S abundance for any experimental group (Figure S7). DNA was 16S rRNA amplicon sequenced, only samples from boxes that passed quality control for both Pre and Post timepoints were used in further analyses to assess the diversity and relative abundance of gut bacteria. Amplicon sequence variants (ASV) were identified and assigned taxa using the BEExact database (Daisley and Reid, 2021). The mean relative abundance of sample genera was plotted per experimental group.

Both experimental groups had comparable mean relative abundances at the Pre timepoint and no sample ASV matched with negative controls. Experimental groups were dominated by five genera *Lactobacillus*, *Gilliamella*, *Bombiscardovia, Bombilactobacillus* and *Snodgrassella* (Figure 6A) which are all core bumblebee gut bacteria (Meeus *et al*., 2015; Kwong *et al*., 2017). Genus *Snodgrassella* was the most abundant occupying ∼50% of experimental group mean relative abundance, with other core taxa occupying varying amounts of the remaining 99% mean relative abundance.

**Figure 6.**
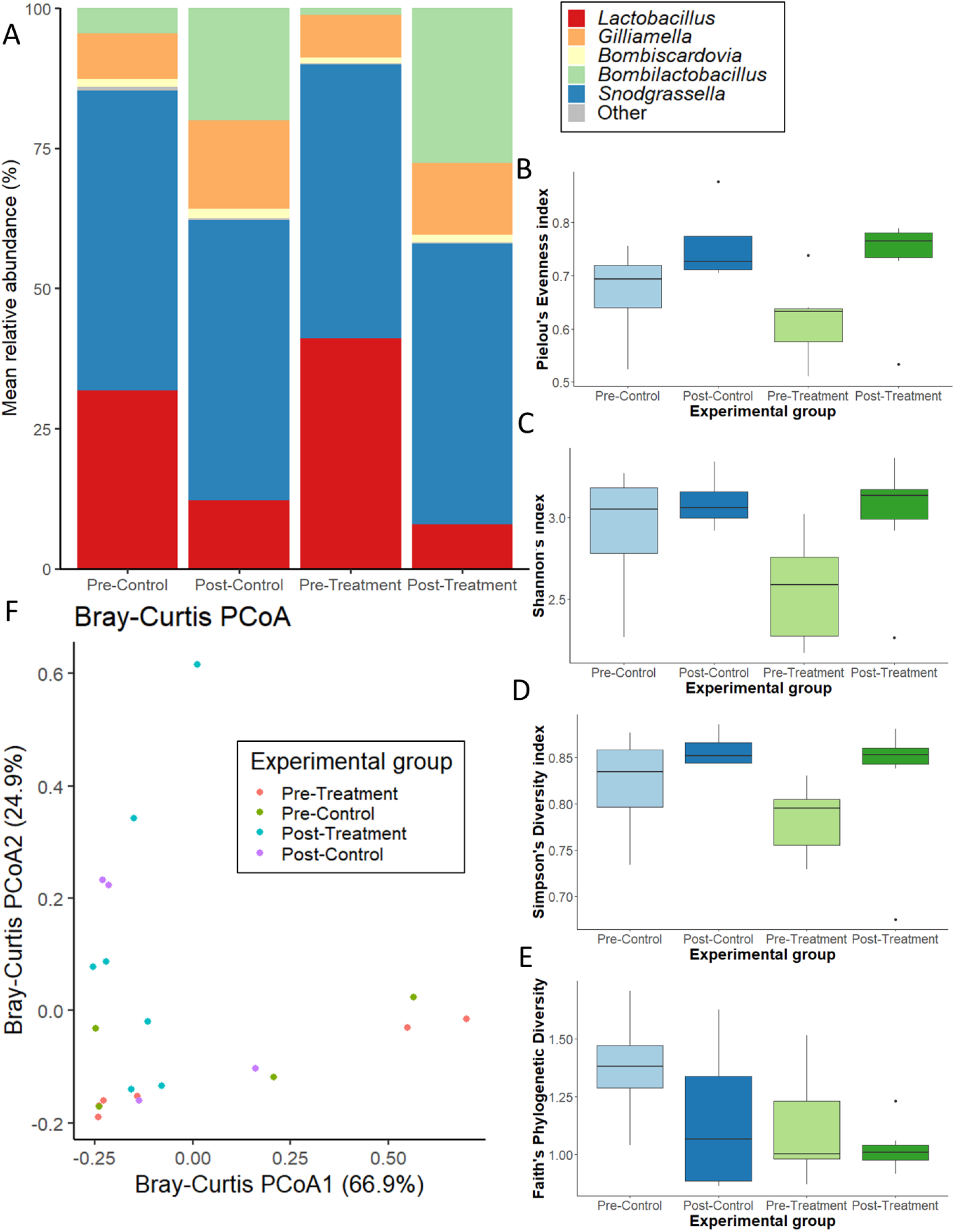
The effect of black carbon on the composition and diversity of adult *Bombus terrestris* gut microbiome. A) Mean relative abundance of genera per experimental group. 16S rRNA amplicon sequences were assigned taxa using the BEExact database in Qiime2 and visualised with the phyloseq package in R. Taxa that were <1% of the total relative abundance are categorised in ‘Other’. Alpha diversity metrics Pielou’s evenness index (B), Shannon’s index (C), Simpsons’s diversity index (D) and Faith’s phylogenetic diversity (E) were calculated for each experimental group. Kruskal-Wallis pairwise tests, and False Discovery Rate corrections were conducted for each alpha diversity metric and found no significant differences at the 0.05 level between control and black carbon treatment groups. F) Bray Curtis PCoA showing microbiome dissimilarity by experimental group, a PERMANOVA calculation found no significant difference between groups at the 0.05 level. PCoA visualised with the phyloseq package in R, axes explain 91.8% of variation.

Alpha and beta diversity analyses were conducted to determine diversity within and between experimental groups. Higher alpha diversity values indicate more diversity and evenly distributed genera. There were increases in the alpha diversity metrics Pielou’s evenness index, Shannon’s index and Simpson’s index, Post black carbon treatment compared to Pre treatment (Figure 6B, 6C, 6D, 6E), yet these differences were not significant (*q* > 0.05). Bray Curtis PCoA plot showed some similarity within experimental groups, particularly clustering Post black carbon treatment, however these differences were not significant with PERMANOVA analyses (*F*_(4, 22)_ = 1.42, *p* > 0.05) (Figure 6F).

The statistical test, analysis of compositions of microbiomes with bias correction (ANCOM-BC) (Lin and Peddada, 2020), was used to identify differentially abundant genera in groups. After black carbon treatment core bacterial taxa *Gilliamella* and *Bombilactobacillus* were significantly differentially abundant (*q* < 0.001) compared to all other experimental groups (Table 1). No other experimental group had any differentially abundant core bacteria. Therefore, black carbon treatment significantly alters the adult *B. terrestris* gut microbiome, changing the abundance of core symbionts *Gilliamella* and *Bombilactobacillus*. These findings provide strong evidence that even at a sublethal dose, exposure to black carbon particulates alter the core gut microbiome of adult *B. terrestris*.

**Table 1.**
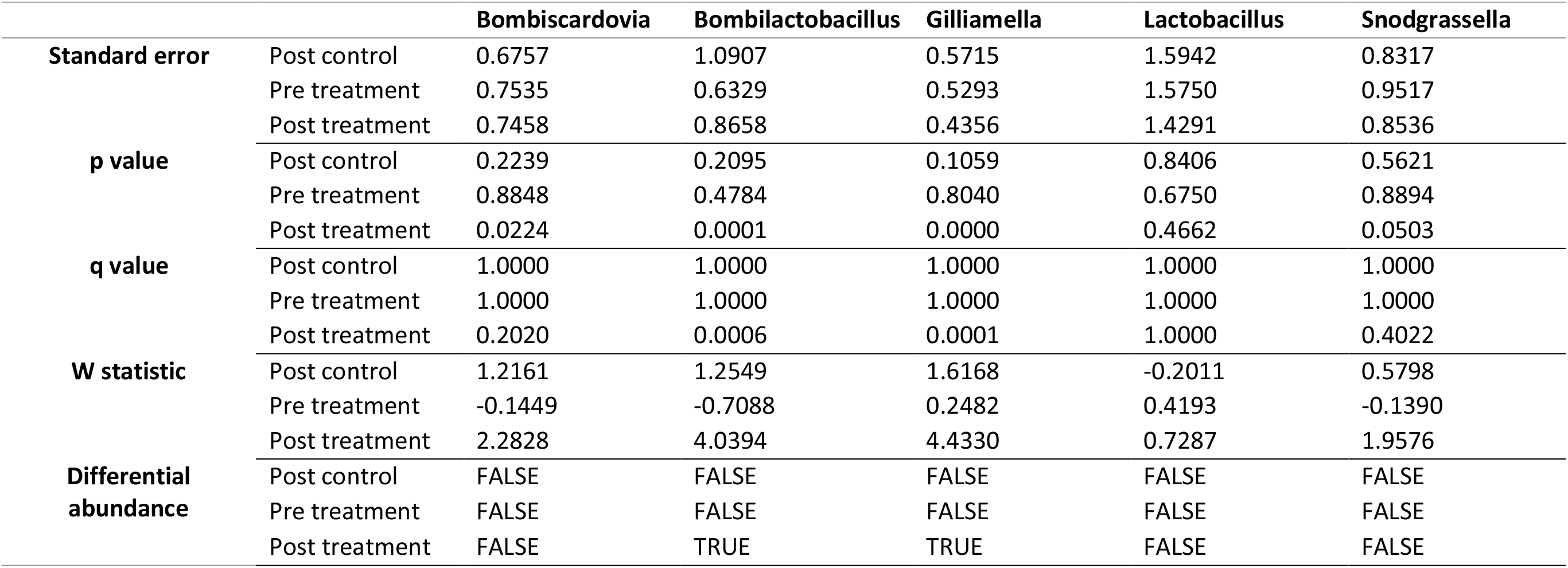
Differential abundance ANCOM-BC scores of core bacterial taxa, analysed by experimental group.

|  |  | **Bombiscardovia** | **Bombilactobacillus** | **Gilliamella** | **Lactobacillus** | **Snodgrassella** |
| --- | --- | --- | --- | --- | --- | --- |
| **Standard error** | Post control | 0.6757 | 1.0907 | 0.5715 | 1.5942 | 0.8317 |
|  | Pre treatment | 0.7535 | 0.6329 | 0.5293 | 1.5750 | 0.9517 |
|  | Post treatment | 0.7458 | 0.8658 | 0.4356 | 1.4291 | 0.8536 |
| **p value** | Post control | 0.2239 | 0.2095 | 0.1059 | 0.8406 | 0.5621 |
|  | Pre treatment | 0.8848 | 0.4784 | 0.8040 | 0.6750 | 0.8894 |
|  | Post treatment | 0.0224 | 0.0001 | 0.0000 | 0.4662 | 0.0503 |
| **q value** | Post control | 1.0000 | 1.0000 | 1.0000 | 1.0000 | 1.0000 |
|  | Pre treatment | 1.0000 | 1.0000 | 1.0000 | 1.0000 | 1.0000 |
|  | Post treatment | 0.2020 | 0.0006 | 0.0001 | 1.0000 | 0.4022 |
| **W statistic** | Post control | 1.2161 | 1.2549 | 1.6168 | -0.2011 | 0.5798 |
|  | Pre treatment | -0.1449 | -0.7088 | 0.2482 | 0.4193 | -0.1390 |
|  | Post treatment | 2.2828 | 4.0394 | 4.4330 | 0.7287 | 1.9576 |
| **Differential abundance** | Post control | FALSE | FALSE | FALSE | FALSE | FALSE |
|  | Pre treatment | FALSE | FALSE | FALSE | FALSE | FALSE |
|  | Post treatment | FALSE | TRUE | TRUE | FALSE | FALSE |

## Discussion

In this study, we have shown that black carbon particulates significantly alter biofilm formation and gene expression of key bee gut commensal, *Snodgrassella alvi* and disrupt adult *Bombus terrestris* gut microbiome composition, increasing viable gut bacteria. Our data provides the first evidence that black carbon alters the behaviour of bees’ commensal bacteria and their gut microbiome composition. Consequently, these findings provide evidence that air pollution directly impacts beneficial commensal bacteria which may contribute to air pollutions’ detrimental effects on bee health.

*S. alvi* is a key, beneficial member of the core gut microbiome in honeybees and bumblebees (Kwong *et al*., 2017) and predominantly colonises the ilea forming a biofilm structure (Martinson *et al*., 2012; Hammer *et al*., 2021). We found that treatment with black carbon changed *S. alvi* wkB2 behaviour, significantly increasing initial adhesion to an abiotic surface *in vitro* and changing early biofilm formation. Our data show an increase in the number of viable biofilm cells at 2, 8 and 24 hours but no change at 48 hours and no overall effect of black carbon on *S. alvi* growth. Black carbon was not incorporated with adhering cells at 2 hours and therefore could not provide a physical support for biofilm formation. Thus, these results suggest that black carbon directly induces *S. alvi* adherence to a surface.

Transcriptional analysis showed that exposure to black carbon significantly decreased the expression of the gene *lysR* in *S. alvi* biofilm cells. As *lysR* is a transcriptional regulator with diverse functions, the change in expression with black carbon treatment likely has further effects on *S. alvi* functions and internal metabolism. LysR homologues in other bacteria, including the closely related species *Neisseria gonorrhoeae*, are involved in the regulation of colonisation and biofilm formation (Seib *et al*., 2007; Maddocks and Oyston, 2008; Apicella *et al*., 2011). The expression of *pilD* in *S. alvi* 24-hour biofilms was also tested, finding no difference in expression with black carbon treatment. As black carbon significantly increases *S. alvi* early biofilm viability, the expression of biofilm related genes *pilD* and *lysR* may also be altered by black carbon in a time related manner. Therefore, while the function of the *S. alvi lysR* gene has not been tested, the significant alteration in *lysR* transcription in black carbon treated *S. alvi* biofilms suggests that *lysR* may play a role in *S. alvi* biofilm regulation. These data provide evidence of a genetic basis for the phenotypic changes of black carbon on *S. alvi* biofilms, consistent with other bacteria that show adaptive genetic responses to black carbon and urban particulate pollutants (Yadav *et al*., 2020; Purves *et al*., 2022).

Scanning electron microscopy images showed that the presence of black carbon dramatically alters the structure of *S. alvi* biofilms. Black carbon particulates were observed incorporated into the 24-hour biofilm structure and there was a significant increase in the size of biofilm protrusions. Significant changes to biofilm protrusion structures were not observed in biofilms exposed to black carbon pre-treated media or chemically inert quartz particles. Particulate pollutants alter biofilm structures in other bacterial species treated with black carbon (Hussey *et al*., 2017) and different particulate pollutants (urban dust and fine particulate matter) (Woo *et al*., 2018; Yadav *et al*., 2020). These studies also found particulate pollutants incorporated into the biofilm (Woo *et al*., 2018; Yadav *et al*., 2020) and showed similar protrusion structures in black carbon treated *S. aureus* (Hussey *et al*., 2017). The mechanisms involved were not established.

We propose that black carbon particulates alter *S. alvi* biofilm formation through physical interaction with the bacteria and induction of a genetic adaptive response. Our data shows that physical interaction between the black carbon particulates and *S. alvi* plays a critical role in biofilm formation because the presence of black carbon particulates is required to change the biofilm structure. There is no change in biofilm formation when *S. alvi* are grown in medium pre-treated with black carbon in the absence of particulates. Notably, the change in biofilm structure is not only through particle-bacteria physical interactions supporting the biofilm structure because *S. alvi* initial adhesion is induced before black carbon particulates are detected in the adherent biofilm cells. Additionally, black carbon alters the expression of the *lysR* gene, a potential regulator of *S. alvi* biofilm formation.

Importantly, increased early biofilm formation along with larger biofilm protrusions of prominent core gut bacteria *S. alvi in vitro* suggests that spatial organisation of microbes in the gut, such as *S. alvi* on the ileal epithelium, could be disrupted with exposure to black carbon *in vivo*. Disrupting the gut microbiome organisation may in turn alter the abundance of other core gut microbes. Indeed our *in vivo* results show longer-term changes to the mature bee gut microbiome with black carbon exposure.

To study the impact of black carbon on *B. terrestris* gut microbiome, age controlled adult worker bees, with a fully formed mature gut microbiome, were subjected to an exposure relevant concentration of black carbon through consumption in their apiary solution. *B. terrestris* exposed to black carbon did not have significantly different survival compared to the control group. Overall, ∼80% of *B. terrestris* from each experimental group survived until the end of the experiment, which would be expected at their age (Smeets and Duchateau, 2003). Additionally, black carbon had no significant effect on *B. terrestris* activity levels compared to controls. Therefore, this concentration of black carbon was not overtly toxic to *B. terrestris* over the experimental period.

There is evidence of the impact of air pollution on wild bee survival showing that air pollution could be having an under-appreciated impact on bee health. Previous work on solitary and social bee species report an association between high environmental air pollution levels and a lower abundance of wild bees (Bromenshenk *et al*., 1991; Moron *et al*., 2012; Thimmegowda *et al*., 2020).

Previous studies have also shown pollution-mediated changes to honeybee foraging behaviour, *A. dorsata* floral visitation rates were significantly decreased when exposed to high air pollution levels (Thimmegowda *et al*., 2020). Furthermore pollution exposure impairs *Apis mellifera* cognitive processes (olfactory learning and memory) that are important for foraging (Leonard, Pettit, *et al*., 2019; Leonard, Vergoz, *et al*., 2019). Bumblebees are primitively eusocial insects and therefore any changes to foraging (and subsequent pollination) would have secondary effects on colony success, pollination rates and ultimately our food production.

Our data demonstrates that although the concentration of black carbon used in these experimental conditions does not overtly affect bee health, black carbon significantly increases viable *B. terrestris* gut bacteria. *B. terrestris* had significantly more MRS culturable bacteria (*Bombilactobacillus* and *Lactobacillus* Firm-5) Post black carbon treatment compared to Pre treatment. Furthermore, 16S rRNA amplicon sequencing of these samples revealed that black carbon significantly alters the composition of *B. terrestris* mature gut microbiome, changing the abundance of beneficial core phylotypes *Gilliamella* and *Bombilactobacillus*. Importantly, *B. terrestris* were exposed to black carbon after obtaining their fully formed, mature gut microbiome, demonstrating that black carbon treatment disrupts the established bee gut microbiome.

Previous work investigating the impact of the heavy metal selenate on *B. impatiens* (Rothman *et al*., 2019) and diesel exhaust particles on *B. terrestris* (Seidenath *et al*., 2023) also saw alterations in the abundance of core gut symbionts in bees treated with these pollutants. In comparison to control bees, both studies found a lower abundance of *Snodgrassella* and *Lactobacillus bombicola* (*Lactobacillus* Firm-5) in pollution treated bees, Seidenath *et al.,* (2023) also saw changes in *Lactobacillus apis* (*Lactobacillus* Firm-5) and *Bombiscardova* abundance with pollution treatment. Changes in these core microbes were not observed in our data. Both previous studies identified changes in *Gilliamella* abundance in pollution treated bees, which agrees with our data, but we discovered changes in *Bombilactobacillus* abundance which was not found in previous work. Consequently, these studies highlight the importance of fully establishing the impact of different pollutants on insect pollinator health because different pollutants have distinct, specific effects on the abundance of core bee gut microbes.

*Bombilactobacillus* (*Lactobacillus* Firm-4) is a core bee gut bacteria that is involved in the metabolism of pollen, short chain fatty acids and glycosides and influences hormonal signalling (Kesnerova *et al*., 2017; Zheng *et al*., 2017). *Gilliamella* is a beneficial core bee gut microbe which is negatively associated with the trypanosome *Crithidia bombi* and is involved in the breakdown of pectin in the pollen cell wall and a range of toxic dietary sugars (Koch and Schmid-Hempel, 2011b; Engel *et al*., 2012; Zheng *et al*., 2016; Kesnerova *et al*., 2017; Zhengyi Zhang *et al*., 2022). Black carbon exposure altered the abundance of these core gut microbes in *B. terrestris*, consequently, this may affect the functionality of the bee gut microbiome such as the ability to utilise a variety of different food sources and to resist pathogen infection. Our findings show that short term ingestion of black carbon has distinct, significant effects on the composition of *B. terrestris* core bacterial community. These results could describe a key mechanism whereby air pollution affects bee physiology and may have serious implications for host health and pollination.

Air pollution particulate matter has a pervasive presence in our atmosphere and accumulates on bee’s bodies and in their food stores. Yet, there is limited knowledge on how particulates affect bee health and to what extent. In this study, we focused our investigations on the core bee gut microbiome as its composition is common to major pollinators (honeybees and bumblebees) and is strongly linked to bee health (Kwong *et al*., 2017; Raymann *et al*., 2017; Rothman *et al*., 2019). We have shown that the major particulate air pollutant, black carbon, has significant effects on core bacteria, differentially altering the composition of *Bombus terrestris* beneficial core gut microbiome. Together our data show that particulate air pollutants are an underexplored risk for the health and balance of environmental microbes and their important natural ecosystems. Policy discussions concerning air pollution and emission reduction would benefit from considering agriculturally important pollinators for the future of our global food security.

## Acknowledgements

This work was supported by a British Beekeepers Association Grant awarded to JAM. HRS was supported by a University of Leicester PhD studentship. We thank Jo Purves for manuscript proofreading, experimental advice and protocols, Alun Jones and Lillie Purser for manuscript proofreading and Novogene for the 16S rRNA library preparation and sequencing.

