## supplementary data for "Air pollution modifies key colonization factors of the beneficial bee gut symbiont *Snodgrassella alvi* and disrupts the bumblebee (*Bombus terrestris*) gut microbiome"

**Table S1. Primers designed (*) and used in this study.**

| Gene name | Direction | Sequence 5' – 3' | Tm (°C) | Product length (bp) |
|---|---|---|---|---|
| LysR* | Forward | TACCCAGACCACTAGCCACC | 60 | 140 |
| | Reverse | CGAAGGCAACTGTATGCGTG | | |
| PilD* | Forward | GAACATGAAGCACAGCGTCC | 60 | 185 |
| | Reverse | ATGATGGAACGAAGCTGGCA | | |
| 16S negative control* | Forward | GCAGCAGTGGGGAATTTTGG | 60 | 138 |
| | Reverse | GTACCGTCAGCACTAGGTGG | | |
| 16S rRNA copy number | Forward (28F) | GAGTTTGATCNTGGCTCAG | 60 | 306 |
| | Reverse (334R) | TGCTGCCTCCCGTAGGAGT | | |

$$\text{BC concentration (mg/g)} = \frac{\text{BC mass administered (0.105 mg)}}{\text{average weight of mouse (35 g)}}$$

$$\text{Feed concentration} = \frac{\text{BC concentration} \ * \ \text{average weight of bee}}{\text{Mass of daily intake of apiary solution per bee}}$$

$$\text{Feed concentration} = \frac{0.003\ \text{mg/g} * 225\ \text{mg}}{1.363\ \text{mg}}$$

$$\text{Feed concentration} = 0.495\ \text{mg/g}$$

**Figure S1. Calculation to determine the black carbon concentration to add to bumblebee feed.** This equation takes into account the black carbon (BC) concentration used in previous published animal experiment, the average weight of adult *Bombus terrestris* and the average amount of apiary solution consumed daily.

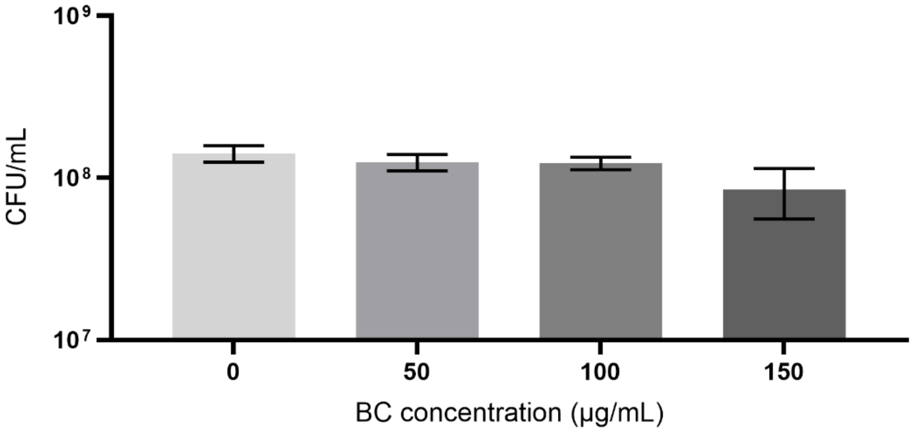


**Figure S2. Black carbon had no significant effect on *S. alvi* growth after 24 h of static incubation.** Overnight cultures of *S. alvi* wkB2 were set to an OD of 0.02 in BHI and supplemented with 0, 50, 100 or 150 µg/mL black carbon (BC). Samples were incubated at 37 °C in 5% $CO_2$ conditions for 24 hours. Serial dilutions were plated onto blood agar and incubated for 48 hours to determine bacterial growth. A one-way ANOVA was performed finding no significant differences in CFU at the 0.05 level, n=3, error bars represent standard error of the mean.

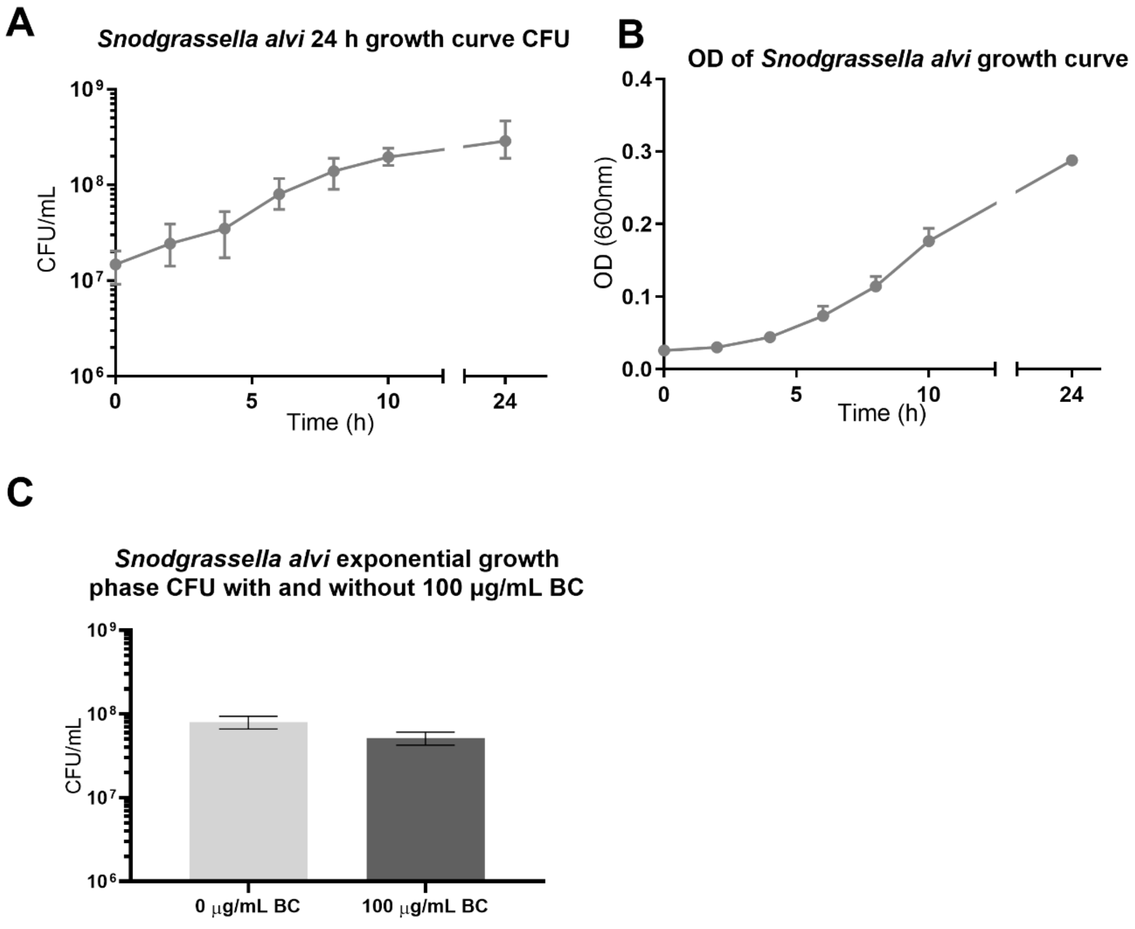

**Figure S3.** ***Snodgrassella alvi*** **growth curve and exponential growth CFU with and without black carbon.** Overnight cultures of *S. alvi* wkB2 were diluted in fresh BHI to an OD of 0.02 and incubated at 37°C in 5% $CO_2$ conditions for 24 hours. At each timepoint samples were taken, serial diluted and plated in technical triplicate to determine CFU (A) and measured for optical density (B). Biological repeats n=4, means of CFU technical triplicates are plotted, error bars represent standard error of the mean. Exponential growth phase of *S. alvi* in these conditions was determined as an $OD_{600\ nm}$ of 0.08 (B). C) Black carbon (BC) had no significant effect on *S. alvi* exponential growth CFU ($t(6) = 1.70$, $p > 0.05$), biological repeats n=4.

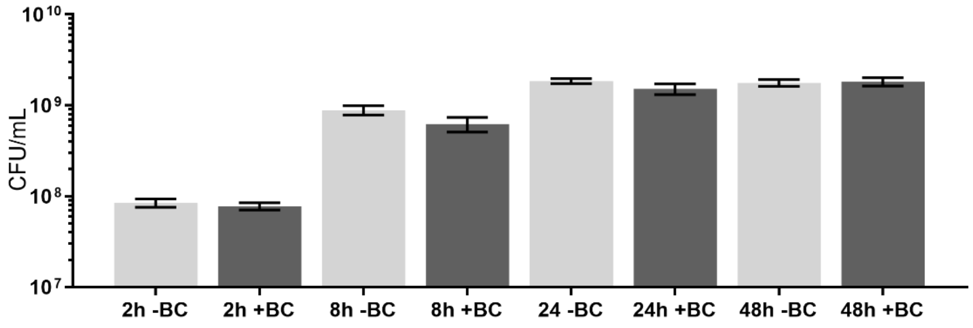


**Figure S4. The effect of black carbon on the total growth of *Snodgrassella alvi* biofilms.** *S. alvi* wkB2 overnight cultures were diluted into fresh BHI with 0 (-BC) or 100 µg/mL black carbon (+BC), aliquoted into 12 well biofilm plates and incubated at 37° C in 5% $CO_2$ conditions for 2 hours, 8 hours, 24 hours or 48 hours. Biofilms were separated into fractions, serial diluted and plated to determine bacterial growth (fraction data presented in Figure 1A, 1B, 1C and 1D). Total CFU for each biofilm timepoint was combined, analysed with a one-way ANOVA and Tukey multiple comparison test finding no significant difference in total CFU with black carbon treatment for any timepoint at the 0.05 level.

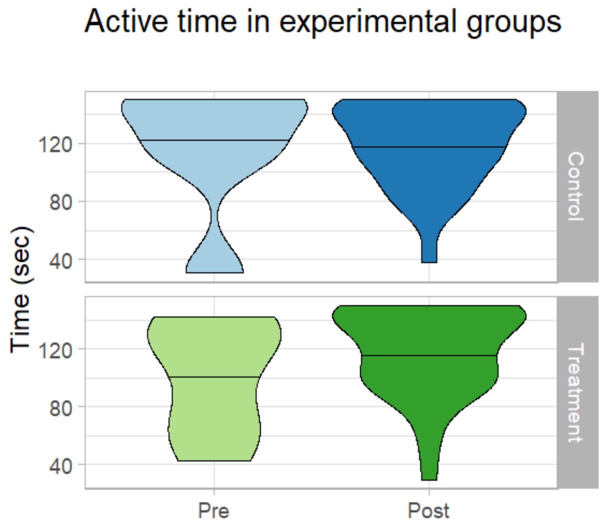


**Figure S5. The effect of black carbon on adult *Bombus terrestris audax* behaviour**. Bees were randomly selected per experimental box and active time was recorded for 150 seconds. This recording took place twice per day and was averaged to a daily mean. Wilcoxon rank sum test with continuity correction was conducted on the daily mean active time within experimental groups finding no significant difference in control or treatment groups, lines represent median.

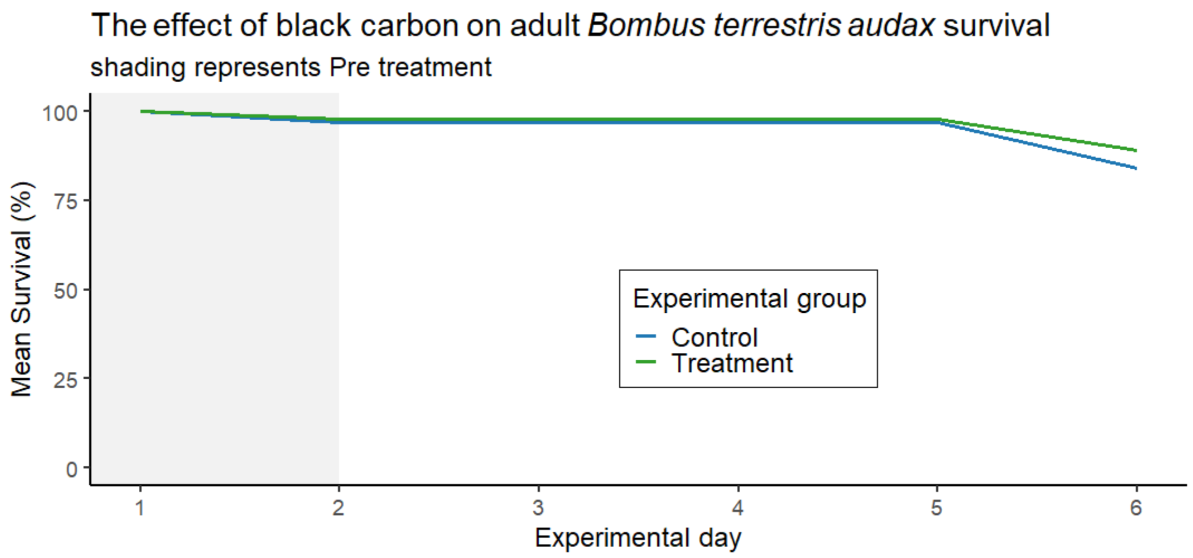


**Figure S6. The effect of black carbon on adult *Bombus terrestris audax* survival**. Survival was recorded per experimental box on days two and six. Percentage survival was determined per box and the mean survival percentage was calculated from these values, finding no significant difference between control and black carbon treatment with the Wilcoxon rank sum test and continuity correction.

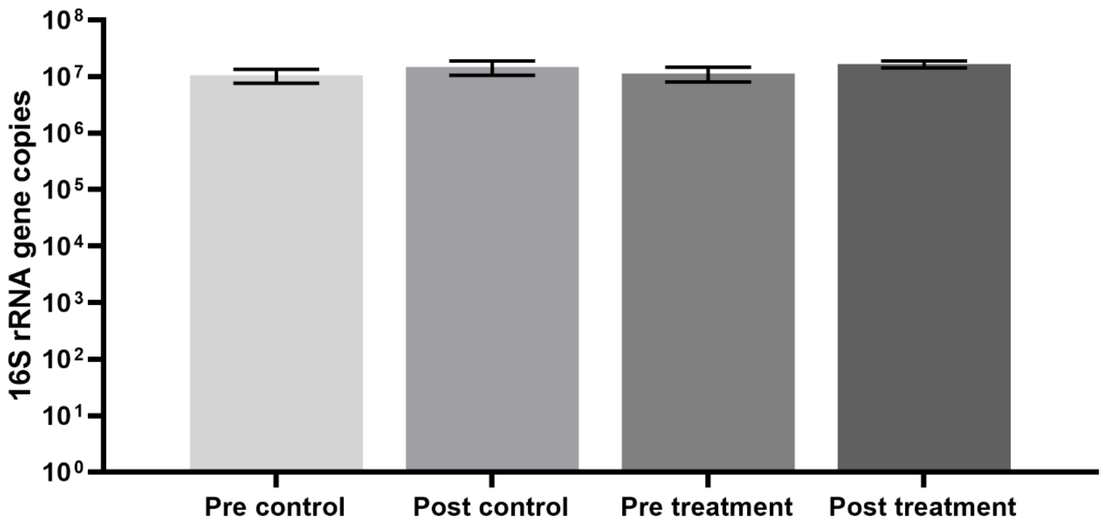


**Figure S7. No significant difference in total bacterial 16S rRNA gene copies between experimental groups.** 16S rRNA copy number was amplified by qPCR in all samples using universal bacterial primers. Copy number per µL of sample was determined using standard curves from the amplification of the cloned target sequence in a pGEM-T vector of known concentration. A one-way ANOVA was performed finding no significant difference between experimental groups at the 0.05 level, error bars represent standard error of the mean.
